# Unified modeling of mutation-induced protein stability responses with OmegaTherm

**DOI:** 10.64898/2026.09.23.753709

**Authors:** Wenkang Wang, Yihang Zhou, Wentao Yu, Yifan Wu, Qiurong Yang, Fanding Xu, Ruiqing Zheng, Xiaoqiang Huang, Min Li, Yang Zhang

## Abstract

Accurate prediction of mutation-induced protein stability responses, including changes in folding free energy and melting temperature, is critical for protein engineering. Existing models represent these related but non-equivalent measurements separately and often focus on single mutations. Here we present OmegaTherm, a sequence-based framework that jointly learns both stability responses for single and multiple mutations. OmegaTherm introduces a shared protein language model to capture transferable mutation context, measurement-specific encoders to preserve measurement identity, and paired-measurement-informed distillation to learn complementary knowledge from heterogeneous data. An antisymmetric prediction architecture further enforces physical consistency between forward and reverse mutations. Across independent benchmarks, OmegaTherm achieves state-of-the-art performance in quantitative prediction, mutation classification and protein-level ranking. Beyond benchmark prediction, OmegaTherm-derived stability landscapes correlate strongly with fitness landscapes from deep mutational scanning, enrich favorable variants and reveal function-associated mutation-sensitive regions. These results establish unified representation learning as an effective strategy for integrating related but non-equivalent biochemical measurements.

## Introduction

Protein stability is a fundamental biophysical property underlying protein folding, function, and evolution, and is central to protein engineering and therapeutic protein development^1,2^. Residue-level mutations can perturb local and long-range interactions, thereby altering protein stability and function^3,4^. Introducing stabilizing mutations is therefore widely used to improve the activity, expression and robustness of engineered proteins^5,6^. Although single mutations provide accessible starting points for mechanistic studies and variant optimization, practical engineering and directed evolution frequently generate variants containing multiple mutations whose combined effects are more difficult to predict. Since the mutational space expands rapidly with mutation number, exhaustive experimental testing is costly and impractical. Accurate modeling of the stability responses induced by single and multiple mutations is needed to uncover sequence–structure– stability relationships that underlie protein behavior and provide a rational foundation for stability-oriented protein engineering^7^.

Mutation-induced stability changes are commonly quantified by the changes in folding free energy (ΔΔG) and melting temperature (ΔT_m_), which characterize thermodynamic and thermal stability responses, respectively. Although these two biophysical measurements are distinct and not interchangeable, both reflect perturbations of the underlying protein folding energy landscape and often exhibit substantial empirical correlation. Both quantities are antisymmetric with respect to mutation reversal: reversing a mutation changes the sign, but not the magnitude, of the stability response. These shared physical properties suggest that joint modeling of ΔΔG and ΔT_m_ can enable complementary knowledge transfer while preserving their distinct biophysical meanings.

Existing methods predict stability changes using different representations of protein sequence and structure. Structure-based approaches, including EvoEF^8,9^, STRUM^10^, and Korpm^11^, use geometric or energy-based descriptions but depend on reliable protein structures. Sequence-based methods such as DDGun^12,13^ and Mutate Everything^14^ avoid this dependency but have more limited access to the residue interactions that determine stability. Protein language models (PLMs)^15–17^ offer an alternative by encoding evolutionary and latent structural information directly from sequence and have recently been adapted to stability prediction^18,19^. However, general PLMs are not specifically optimized for mutation-induced perturbations, particularly when modeling multiple mutations. These requirements thus motivate mutation-centered modeling that explicitly contrasts wild-type and mutant states, organizes sequence and contact information around the mutated residues, and aggregates perturbations across one or multiple mutation sites.

Beyond the representation of mutation effects, the fragmented and heterogeneous nature of available experimental stability data poses challenges to unified modeling. ΔΔG and ΔT_m_ datasets differ markedly in scale, experimental conditions and label distributions, with only limited overlap in experimentally paired mutations. Existing predictors therefore typically treat the two measurements as separate learning problems, limiting their ability to exploit shared stability-relevant information. Directly pooling the datasets is inappropriate because ΔΔG and ΔT_m_ retain distinct biophysical meanings, whereas unconstrained cross-measurement transfer may propagate unreliable supervision and measurement-specific biases. Unified modeling therefore requires reliable knowledge transfer across heterogeneous and largely unpaired datasets without propagating unreliable supervision or obscuring measurement-specific information.

Here we present OmegaTherm, a sequence-based framework for unified modeling of mutation-induced protein stability responses. Through mutation-supervised finetuning, OmegaTherm first adapts the shared PLM encoder to generate mutation-responsive sequence representations and corresponding residue contact information. Measurement-specific mutation-centered encoders then integrate these shared features around mutation sites while retaining distinct response patterns for ΔΔG and ΔT_m_. To learn from heterogeneous and incompletely paired stability data, self-and cross-measurement distillation further transfer complementary supervision while filtering inconsistent pseudo labels. An antisymmetric architecture additionally enforces sign reversal between forward and reverse mutations. Across comprehensive benchmarks, OmegaTherm achieved state-of-the-art performance across both stability measurements and mutation complexity. Its predictions were also associated with DMS-derived fitness landscapes and enriched high-fitness variants, while downstream analyses highlighted mutation-sensitive regions. Together, these results show that OmegaTherm can unify complementary measurements of protein stability and support rational protein engineering.

## Results

### OmegaTherm unifies mutation-induced stability modeling across measurements and mutation complexity

OmegaTherm was designed to unify the modeling of mutation-induced thermodynamic (ΔΔG) and thermal (ΔT_m_) stability responses across single and multiple mutations. Its unified design couples a shared PLM encoder with measurement-specific mutation-centered transformer encoders, allowing both measurements to benefit from common mutation-responsive sequence representations while retaining their distinct features and prediction outputs.

To adapt pretrained knowledge to the physical consequences of mutations, OmegaTherm separately encodes wild-type and mutant sequences to obtain residue representations and sequence-derived contact information (Fig. 1a). For each mutation site, it constructs paired features that combine wild-type and mutant representations, together with their differences and element-wise interactions, while retaining both sequentially adjacent and contact-defined residues (Fig. 1b). Measurement-specific mutation-centered encoders subsequently integrate these local and long-range contexts across mutation sites, producing measurement-specific representations that accommodate both single and multiple mutations (Fig. 1c). The resulting representations are finally combined with measurement-relevant experimental conditions (e.g., pH and temperature for ΔΔG, and pH for ΔT_m_) and processed by an antisymmetric architecture that enforces the physical sign-reversal constraint between forward and reverse mutations (Fig. 1d).

**Fig. 1.**
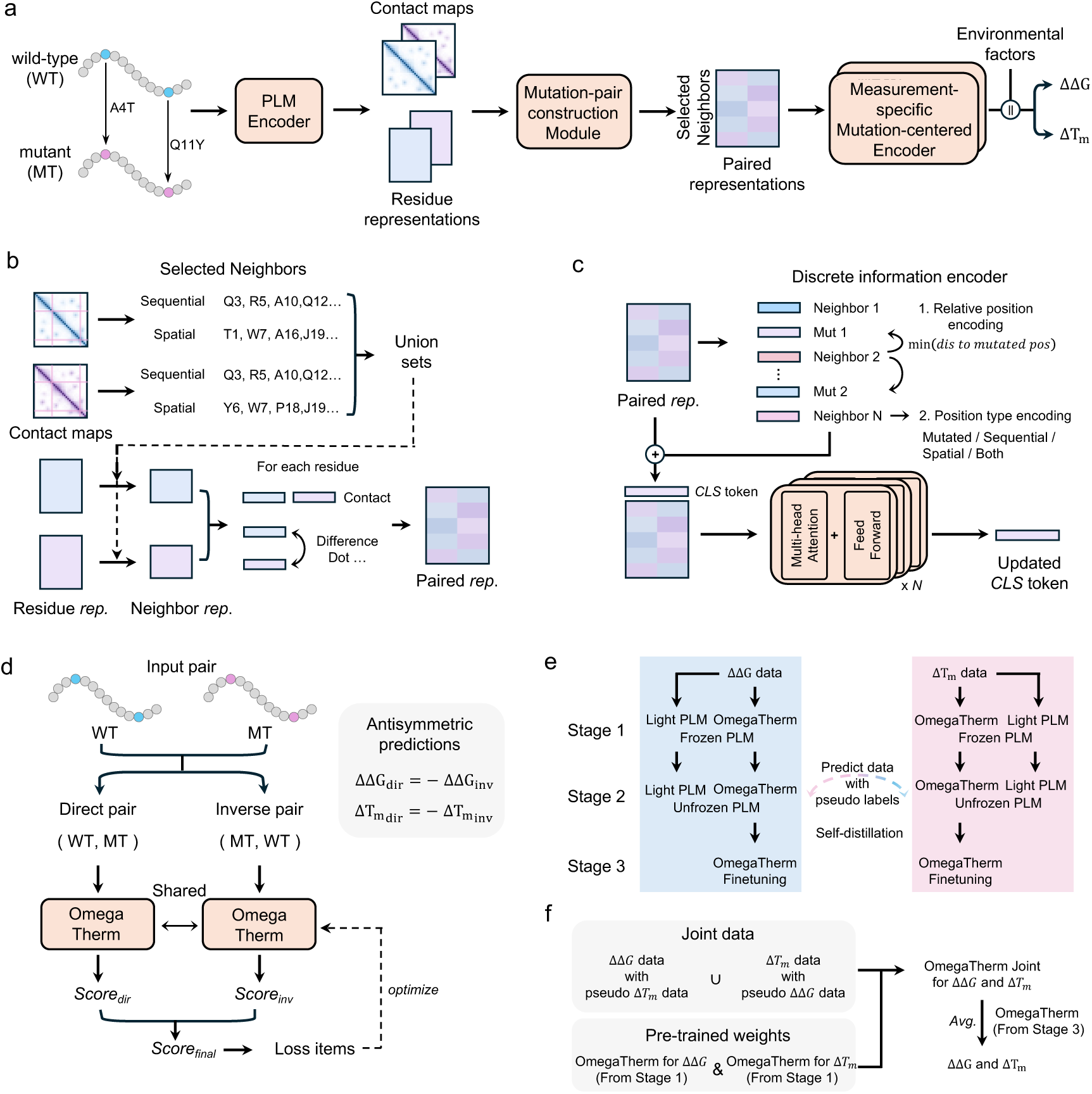
Model architecture of OmegaTherm. **a**, Overview of OmegaTherm. Wild-type (WT) and mutant (MT) protein sequences are separately encoded to generate sequence representations and predicted contact maps. These features are integrated into mutation pair-level representations, processed by a mutation-centered encoder and combined with environmental parameters, including pH and temperature, to predict stability changes. **b**, Mutation-pair construction module. **c**, Mutation-centered transformer encoder. **d**, OmegaTherm training workflow. WT and MT sequences are used to construct direct and inverse pairs for input into the model. **e**, Self-distillation strategy. **f**, Cross-distillation strategy for unified model construction.

To learn from heterogeneous and largely unpaired measurements, OmegaTherm further employs self-and cross-measurement guided by the physical consistency constraints. Task-specific teacher models first generate complementary pseudo labels for unpaired stability responses across ΔΔG and ΔT_m_, thereby increasing supervision beyond the limited experimentally paired measurements. Pseudo labels are filtered by excluding inconsistent signs or implausible magnitudes. The remained high-quality pseudo labels are used both to iteratively refine the task-specific models through self-distillation (Fig. 1e) and to train a unified student through cross-distillation (Fig. 1f). This distillation strategy enables knowledge transfer between thermodynamic and thermal stability measurements while limiting the propagation of unreliable pseudo labels.

### OmegaTherm accurately predicts, classifies, and ranks thermodynamic stability changes across mutation complexity

We first evaluated OmegaTherm for mutation-induced thermodynamic stability changes using the independent single and multiple mutation benchmarks S461^11^ and M223. Performance was assessed from three complementary perspectives: quantitative prediction of ΔΔG, classification of mutations as destabilizing, neutral, or stabilizing, and protein-level ranking of mutation-induced stability changes.

For quantitative ΔΔG prediction, OmegaTherm achieved the best overall performance on both single and multiple mutations. For single mutations, OmegaTherm achieved the highest SCC (Spearman correlation coefficient) 0.85, PCC (Pearson correlation coefficient) 0.83, and R2 (Coefficient of determination) 0.68, while reducing MedAE (Median absolute error) by 18% relative to the strongest competing method (0.424 vs. 0.512 kcal/mol; Fig. 2a and Supplementary Tables 1 and 2)^12,13,18,20–34^. The performance advantage became more evident for multiple mutations, with improvements of at least 24%, 15%, and 60% in SCC, PCC, and R^2^, respectively, together with reductions of 18% in RMSE (2.07 kcal/mol) and 42% in MAE (1.16 kcal/mol) (Fig. 2a and Supplementary Table S3). Overall, among methods applicable to both single and multiple mutations, OmegaTherm consistently achieved the best performance across all evaluation metrics (Fig. 2b). These results show that OmegaTherm accurately captures thermodynamic stability responses across both individual and combinatorial mutations.

**Fig. 2.**
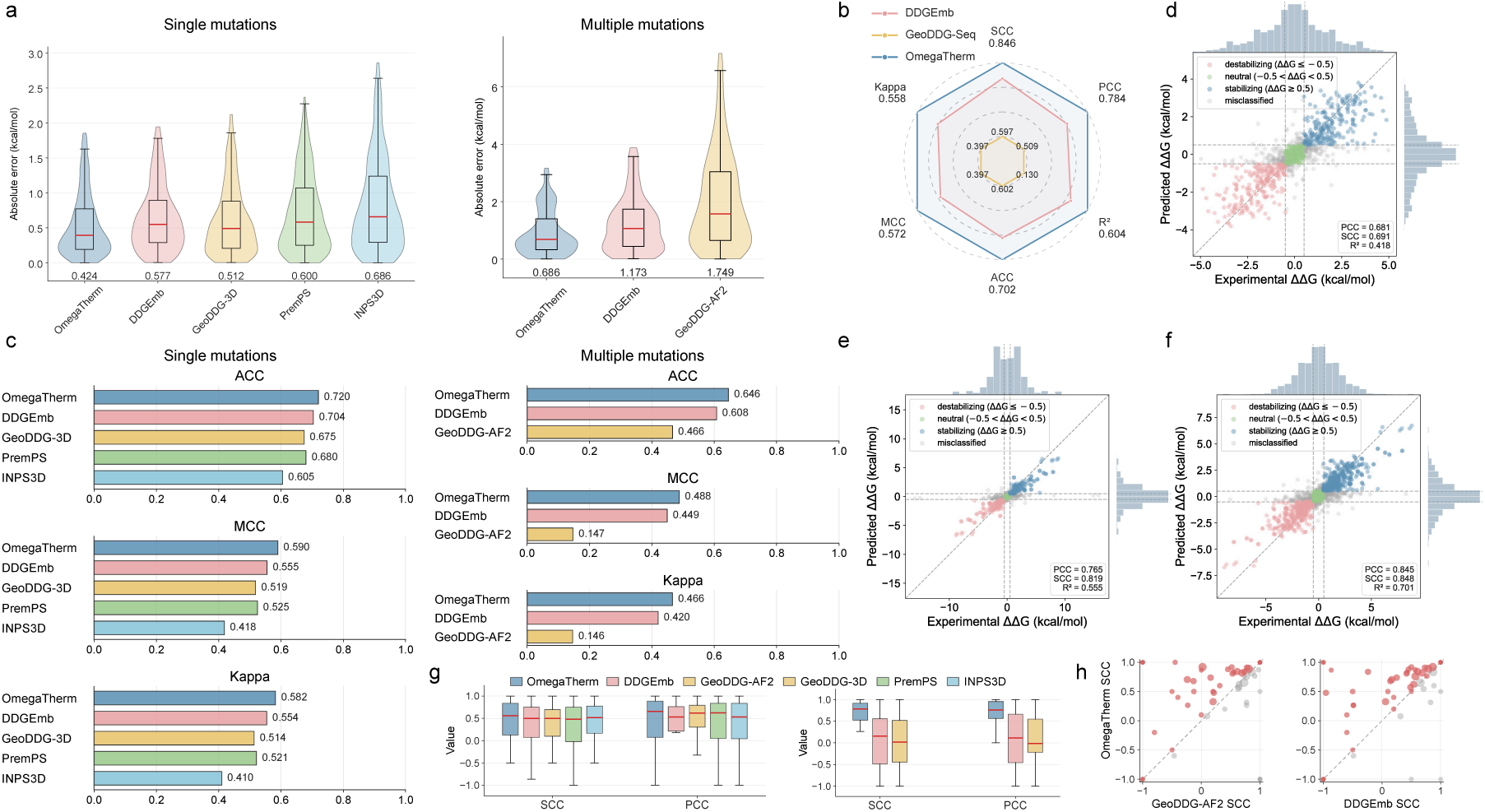
Performance comparison of OmegaTherm for ΔΔ*G* prediction. **a**, Violin plots of AE for OmegaTherm and representative competing methods. Red lines denote MedAE, box bounds indicate the first and third quartiles, and whiskers indicate 0.8 times the interquartile range (IQR). **b**, Performance comparison on both single and multiple mutations. **c**, Scatter plot of OmegaTherm predictions on single-mutation data, with mutations grouped into three classes according to ΔΔG thresholds. **d**, Comparison of ACC, MCC and Cohen’s κ score for ΔΔG classification. **e**, Scatter plot of OmegaTherm predictions on multiple-mutation data. **f**, Scatter plot of OmegaTherm predictions on both single and multiple mutation data after outlier removal. **g**, Protein-level performance comparison between OmegaTherm and other methods, including single mutations (left panel) and multiple mutations (right panel). **h**, Detailed protein-level comparison among OmegaTherm, GeoDDG and DDGEmb, where the size of each point represents the number of mutations present in the protein.

In practical protein engineering cases, distinguishing stabilizing mutations from neutral and destabilizing mutations is crucial^35–37^. We therefore further evaluated whether OmegaTherm by recovering the experimentally defined stability categories (ΔΔG < −0.5 kcal/mol), neutral (|ΔΔG| ≤ 0.5 kcal/mol), and stabilizing (ΔΔG > 0.5 kcal/mol). OmegaTherm achieved the best overall classification performance for both mutation settings (Fig. 2c). It correctly classified 664 of 922 single mutations (Fig. 2d), with both MCC and Kappa exceeding 0.58, compared with approximately 0.55 for the second-best method, DDGEmb. For multiple mutations, OmegaTherm maintained the highest classification performance, with an ACC (0.646) improvement of at least 6% over competing methods (Fig. 2e). Additionally, we further examined the sensitivity of model performance to extreme values (ΔΔG < 10 kcal/mol) by repeating both quantitative and classification evaluations after excluding these mutations. The improved correlation-based performance and unchanged classification accuracy indicate that extreme ΔΔG responses primarily challenge precise magnitude estimation, while categorical discrimination remains robust to their presence (Fig. 2b,f).

Since protein engineering commonly requires prioritizing mutations within the same protein for experimental screening, we next evaluated OmegaTherm by ranking mutation effects within individual proteins. OmegaTherm achieved the highest protein-level SCC and PCC on both benchmarks (Fig. 2g,h). For single mutations, it achieved a median SCC of 0.559 and a median PCC of 0.656, exceeding the best competing methods (0.515 for INPS3D and 0.625 for PremPS, respectively). The advantage was larger for multiple mutations, where OmegaTherm achieved an SCC of 0.783 and a PCC of 0.761, substantially outperformed the strongest competing method (SCC = 0.155; PCC = 0.114) (Fig. 2g,h).

Collectively, these results demonstrate that OmegaTherm showed consistent performance in quantitative prediction, categorical identification, and ranking of ΔΔG responses across single and multiple mutations, thereby providing a reliable computational framework for stability-guided protein engineering.

### OmegaTherm extends unified stability modeling to thermal stability responses

Having established OmegaTherm’s performance for thermodynamic stability responses, we next investigated whether the unified framework extends to mutation-induced thermal stability changes measured byΔT_m_. We benchmarked OmegaTherm on the independent single and multiple mutation benchmarks S571^18^ and M286 using the same three tasks as in thermodynamics evaluation.

For quantitative ΔT_m_prediction, OmegaTherm achieved the strongest overall performance across both single and multiple mutations^18,38–40^. For single mutations, OmegaTherm achieved the lowest RMSE (7.41 °C), MAE (4.89 °C) and MedAE (2.99 °C), while reaching the highest PCC of 0.64, SCC of 0.60, and R^2^ of 0.39, improving these metrics by 26%, 14%, and 67%, respectively, over the strongest competing method (Fig. 3a and Supplementary Table 4). For multiple mutations, OmegaTherm performed best across nearly all metrics (Supplementary Tables 5 and 6), reducing MedAE from 8.318 °C to 7.237 °C (Fig. 3a). Across both mutation settings, it achieved the highest overall SCC of 0.489 (Fig. 3b). Thus, the predictive performance observed for ΔΔG extends to the distinct thermal stability response measured by ΔT_m_.

**Fig. 3.**
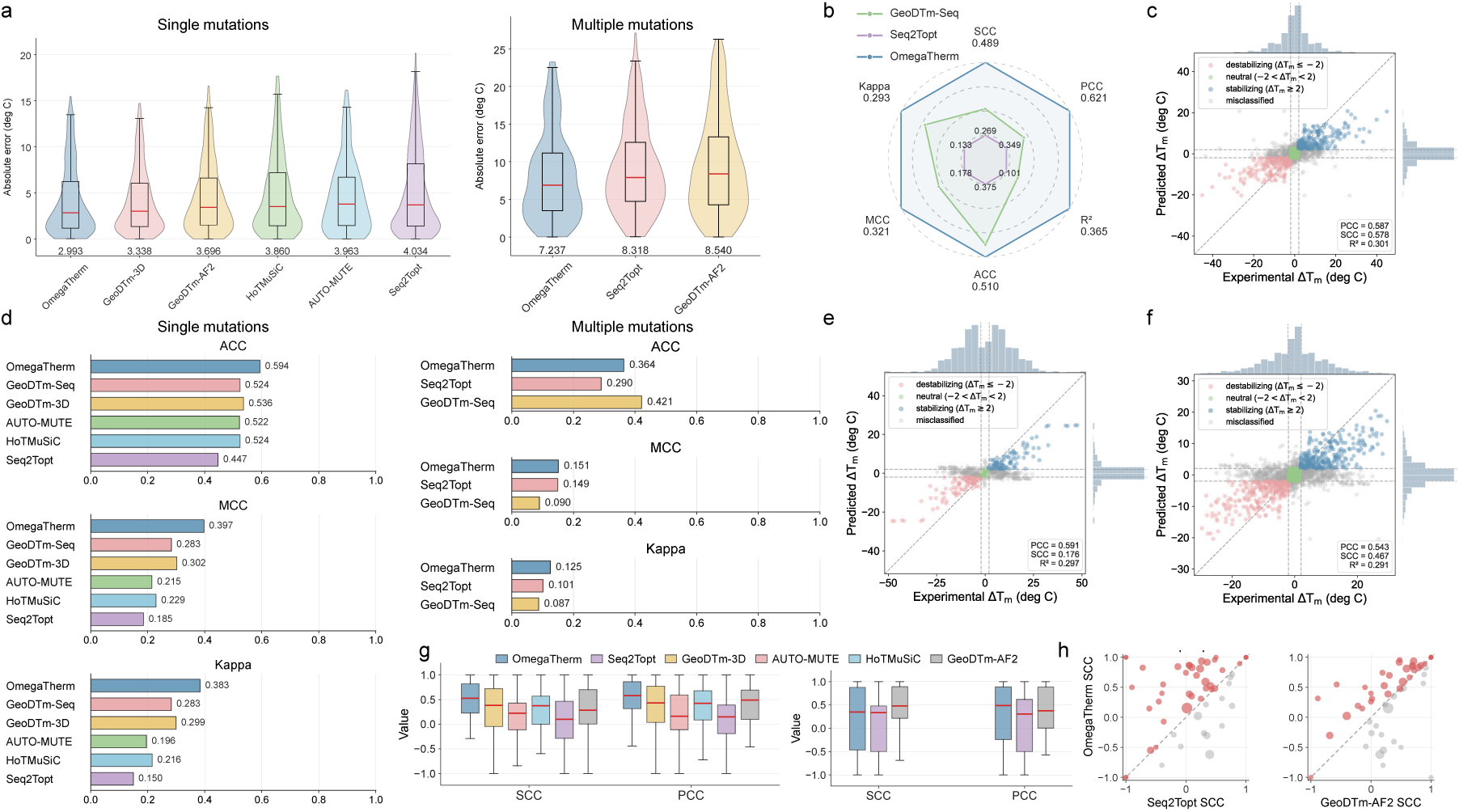
Performance comparison of OmegaTherm for ΔT_m_prediction. **a**, Violin plots of AE for OmegaTherm and representative competing methods. Red lines denote MedAE, box bounds indicate the first and third quartiles, and whiskers indicate 0.8 times the interquartile range (IQR). **b**, Performance comparison on both single and multiple mutations. **c**, Scatter plot of OmegaTherm predictions on single-mutation data, with mutations grouped into three classes according to ΔT_m_thresholds. **d**, Comparison of ACC, MCC and Cohen’s κ score for ΔT_m_ classification. **e**, Scatter plot of OmegaTherm predictions on multiple-mutation data. **f**, Scatter plot of OmegaTherm predictions on both single and multiple mutation data after outlier removal. **g**, Protein-level performance comparison between OmegaTherm and other methods, including single mutations (left panel) and multiple mutations (right panel). **h**, Detailed protein-level comparison among OmegaTherm, GeoDTm and Seq2Topt, where the size of each point represents the number of mutations present in the protein.

The performance advantage extended to mutation classification. OmegaTherm improved classification accuracy for single mutations by 10.8% over the strongest competing method (Fig. 3c,d), while achieving the best performance across most metrics for multiple mutations, correctly classifying 208 of 572 variants (Fig. 3e). Unlike for ΔΔG, excluding extreme ΔT_m_ values reduced PCC, SCC and R², whereas classification performance remained largely unchanged (Fig. 3b,f). These contrasting patterns indicate that extreme responses affect correlation-based evaluation differently for ΔΔG and ΔT_m_, whereas categorical discrimination remains comparatively insensitive to them.

OmegaTherm further maintained strong protein-level ranking performance for ΔT_m_. For single mutations, it achieved the highest median SCC (0.53) and PCC (0.58), representing 36.9% and 33.9% improvements over GeoDTm-3D (Fig. 3g). For multiple mutations, OmegaTherm maintained the highest median PCC (0.49) and a comparable median SCC (0.35) to the best-performing method, showing superior ranking performance across most individual proteins (Fig. 3g,h). These results showed that OmegaTherm could rank mutation-induced thermal stability response across both single and multiple mutations, extending its protein-level ranking capability beyond thermodynamic stability response.

Together, the ΔΔG and ΔT_m_ benchmarks demonstrate that OmegaTherm consistently predicts, classifies and ranks mutation responses across distinct measurements and mutation complexity.

### OmegaTherm generalizes stability modeling to mutation fitness landscapes

To investigate whether the learned stability representations capture biologically meaningful mutation effects beyond stability prediction, we evaluated OmegaTherm on a large deep mutational scanning (DMS) dataset, in which experimental measurements directly quantify the fitness of individual mutations rather than ΔΔG or ΔT_m_^41,42^. We thus used rank-based correlations to assess whether the reconstructed stability landscapes were consistent with experimentally observed fitness landscapes^43^.

Across the complete benchmark, both OmegaTherm-predicted ΔΔG and ΔT_m_ showed strong agreement with experimental mutation fitness, without a marked bias toward either stability measurement (Supplementary Fig. 1). Compared with GeoDDG, OmegaTherm substantially improved SCC and PCC by 13.8% and 11.6% between ΔΔG and experimental fitness across the entire dataset (SCC = 0.643 vs. 0.565; PCC = 0.673 vs. 0.603) as well as at the individual-protein level (median SCC = 0.72 vs. 0.64; median PCC = 0.76 vs. 0.68; Fig. 4a, left). Predictions from the ΔT_m_ head exhibited similarly strong correlations with fitness and substantially outperformed Seq2Topt (Fig. 4a, right). Across individual proteins, OmegaTherm consistently exceeded the performance of both GeoDDG and Seq2Topt in recovering mutation-dependent fitness landscapes (Fig. 4b). These results indicate that the unified model captures a consistent stability-related component of mutation fitness across the two distinct stability dimensions.

**Fig. 4.**
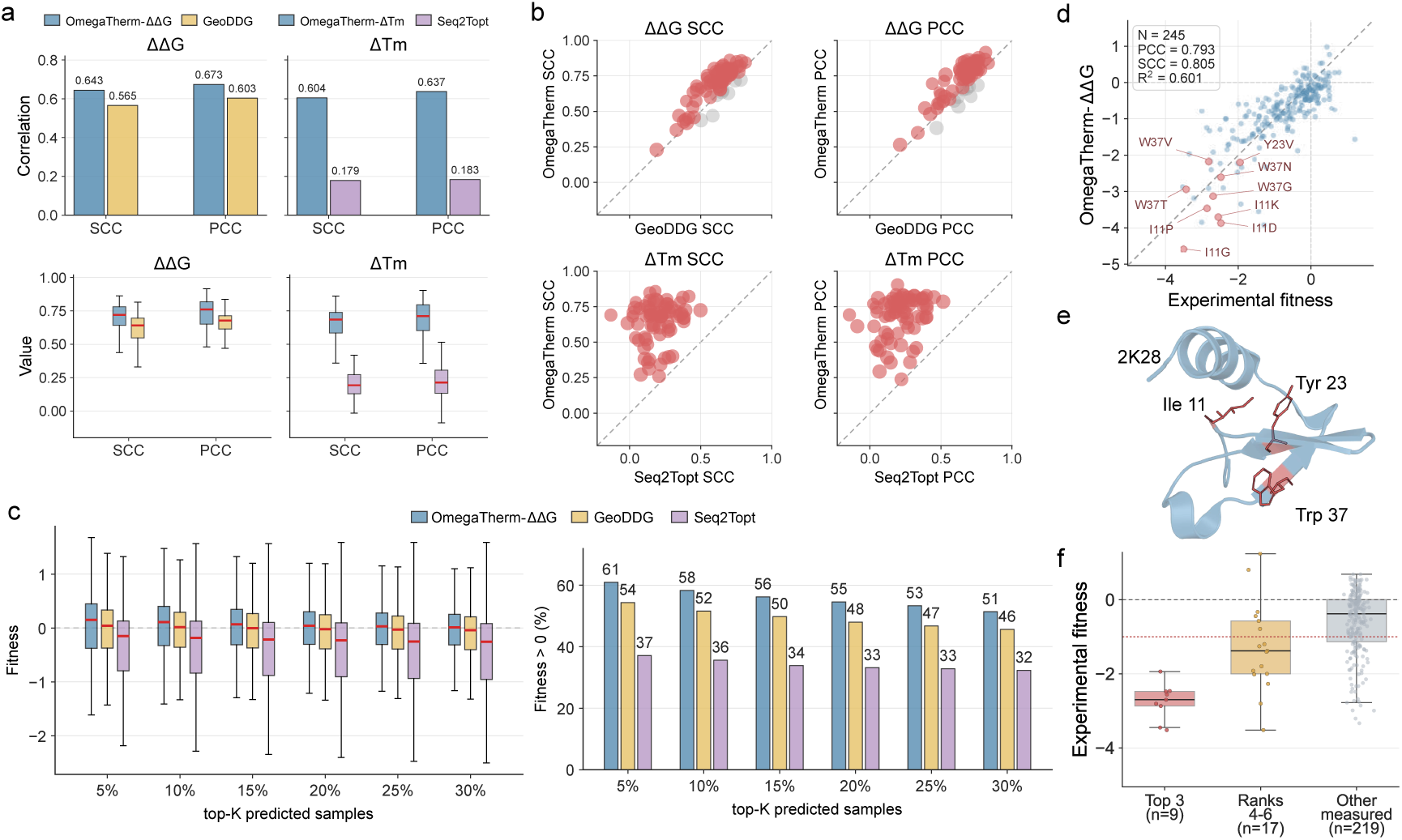
Benchmarking on large-scale DMS data. **a**, Overall performance comparison between OmegaTherm and other tools on 20,000 DMS mutations. The upper panel shows different experimental measurement-level results, and the lower panel shows corresponding protein-level results. **b**, Protein-level comparison of OmegaTherm and other methods using SCC and PCC. Each point in the scatter plot represents one protein with multiple associated mutations. **c**, Left panel is box plots of experimentally measured fitness values for mutations selected at different top-K prediction proportions. Higher fitness values indicate stronger enrichment of stabilizing mutations among top-ranked predictions. Right panel is the proportion of stabilizing mutations in the top-K predictions. **d**, The performance of the OmegaTherm landscape on 2K28. **e**, Top three mutation-sensitive sites in 2K28. **f,** Distribution of fitness changes corresponding to mutations at different sensitivity intervals.

We next extended the ranking analysis from the preceding benchmarks to the more practical task of selecting high-fitness mutations. Mutations were ranked by their predicted ΔΔG or ΔT_m_ values, and the experimentally measured fitness of the top-ranked variants was evaluated at progressively varying top-K fractions. Across all thresholds, OmegaTherm consistently enriched mutations with higher experimental fitness than those selected by GeoDDG or Seq2Topt (Fig. 4c, left). Moreover, OmegaTherm recovered a consistently higher fraction of favorable mutations (fitness > 0) within the same top-ranked candidates (Fig. 4c, right), demonstrating that the learned stability representation more effectively enriches functionally beneficial variants.

Beyond mutation prioritization, the stability landscapes reconstructed by OmegaTherm revealed mutation-sensitive regions that were strongly enriched for annotated functional residues. As an example, OmegaTherm accurately reproduced the experimentally determined mutational landscape of E3 SUMO-protein ligase CBX4 (Fig. 4d, PDB: 2K28) from the DMS dataset and further inferred the complete single-residue substitution landscape (Supplementary Fig. 2, left). Across 50 measured positions, the predicted sensitive scores correlated strongly with the observed median destabilization effects (SCC = 0.786, Supplementary Fig. 3). All nine measured substitutions at the three highest-ranked sites caused strong destabilization (Fig. 4d,e), whereas such effects occurred less frequently at lower-ranked sites (Fig. 4f). Notably, among these three positions, Ile11 corresponded to a previously reported mutagenesis-associated residue whose substitution disrupts interactions with H3C15, H3C1, and RNF246^44^, supporting the biological relevance of the reconstructed stability landscape.

### Ablations identify the contributions of OmegaTherm design components

We next quantified the contributions of three major design components: sequence-derived structural context, mutation-centered adaptation of pretrained representations, and unified learning across heterogeneous stability measurements with distillation.

Although OmegaTherm operates directly from protein sequences, it incorporates sequence-derived residue-contact information inferred by a pretrained PLM model to provide structural context for mutation-effect prediction. Removing contact information consistently reduced prediction accuracy for both ΔΔG and ΔT_m_ prediction. For single mutation prediction, RMSE increased by 4.1% and 4.6% (from 0.964 to 1.004 kcal/mol for ΔΔG and from 7.327 °C to 7.666 °C for ΔT_m_), whereas SCC decreased by 3.2% and 3.7% (0.69 vs. 0.67 for ΔΔG, 0.59 vs. 0.57 for ΔT_m_, Fig. 5a). Similar performance reductions were also observed for multiple mutations (Supplementary Figs. 4 and 5). These results demonstrate that sequence-derived structural context contributes substantially to the learned stability representation.

**Fig. 5.**
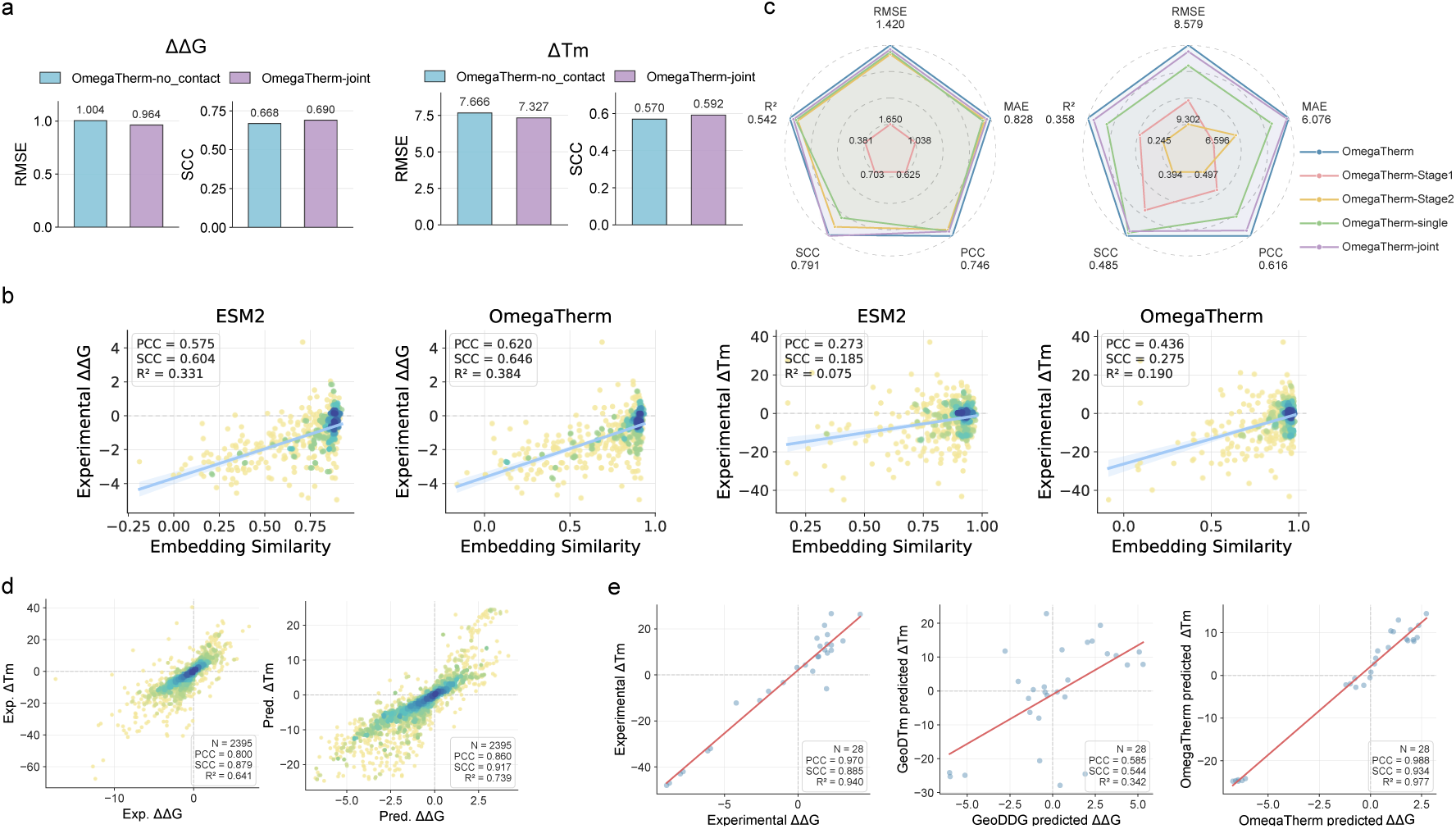
Analysis of the core design principles of OmegaTherm. **a,** Ablation study of structural contact map component in ΔΔG and ΔT_m_ prediction, where OmegaTherm-no_contact denotes the student model without structural neighbors, and OmegaTherm-joint is the final student model. **b,** Correlation between experimental ΔΔG/ΔT_m_ values and model embeddings from the fixed ESM2 model and OmegaTherm. **c,** Performance comparison of different distillation strategies on ΔΔG and ΔT_m_datasets. **d,** Correlations between experimentally verified paired ΔΔG and ΔT_m_ training data and OmegaTherm predictions. **e,** Comparison of correlations in paired ΔΔG and ΔT_m_ test data, where the left panel shows the correlations of the experimental data from the test set; the middle panel shows the correlations of the GeoDDG and GeoDTm prediction results; and the right panel shows the correlations of the OmegaTherm prediction results.

Mutation-supervised adaptation is also important for transferring general protein representations to quantitative stability responses. Finetuning the pretrained PLM consistently outperformed the PLM-frozen version and increased the sensitivity of learned features to residue substitutions (Fig. 5b and Supplementary Fig. 6). This improvement reflect the mismatch between sequence-level pretraining, which emphasizes evolutionary regularities, and mutation-effect prediction, which requires sensitivity to subtle residue perturbations. The effect was particularly pronounced for ΔT_m_, whose PCC, SCC, and R^2^ increased by 59.7%, 48.6%, and 153.3%, respectively (Fig. 5b, right). These findings indicate that mutation-specific supervision effectively adapts general protein representations to mutation-centered stability responses.

We also investigated whether knowledge distillation improves representation learning under limited experimental supervision. During distillation, task-specific teacher models generated complementary pseudo labels that enrich supervision for the unified student model. Removing distillation consistently caused performance deduction across all evaluated metrics (Fig. 5c). For ΔΔG prediction, both OmegaTherm-single and OmegaTherm-joint outperformed OmegaTherm-stage2 across RMSE, MAE, PCC, SCC, and R² (Supplementary Fig. 7). Similar trends were observed for ΔT_m_prediction (Supplementary Fig. 8), demonstrating that cross-distillation improve predictive performance when experimental annotations are limited or unbalanced.

Finally, we evaluated the contribution of unified learning from heterogeneous stability measurements. Jointly trained models consistently performed comparably to or better than their corresponding single-task models (Supplementary Fig. 7 and 8), with the largest improvement observed for ΔT_m_ . This suggests that thermal stability prediction benefits substantially from complementary information contained in ΔΔG measurements. Consistent with this observation, experimentally paired ΔΔG and ΔT_m_ values exhibited a clear positive correlation, and OmegaTherm successfully reproduced this relationship following unified training (Fig. 5d). Furthermore, evaluation on 28 paired mutations from the held-out test set showed that OmegaTherm better reproduced the observed association between ΔΔG and ΔT_m_ than independently trained GeoDDG and GeoDTm models (Fig. 5e). These results support cross-measurement transfer through a shared encoder while preserving separate measurement-specific outputs.

Collectively, these analyses identify complementary contributions from the major OmegaTherm components. Sequence-derived structural context improves mutation-effect prediction without requiring explicit protein structures. Mutation-supervised finetuning adapts general PLMs to mutation-sensitive representations. Knowledge distillation enriches supervision under limited experimental data, and unified learning enables knowledge transfer across heterogeneous stability measurements. Together, these components enable OmegaTherm to generalize across thermodynamic and thermal stability measurements, mutation complexities, and diverse protein families.

## Discussion

OmegaTherm provides strong evidence that related but non-equivalent measurements can be effectively exploited even when observations are heterogeneous and largely unpaired. The framework shares mutation-responsive representations across ΔΔG and ΔT_m_ while retaining measurement-specific encoders for preserving their distinct characteristics. Self-and cross-measurement distillation further transfers complementary knowledge with paired-measurement-informed filtering while an antisymmetric architecture enforces sign reversal between forward and reverse mutations. The resulting performance supports the value of formulating stability prediction from the underlying physical relationships rather than treating individual measurements and mutation settings as independent learning problems.

Across independent benchmarks, OmegaTherm achieved state-of-the-art performance for both ΔΔG and ΔT_m_ across single and multiple mutations, while maintaining consistent capability across the two stability dimensions in quantitative prediction, mutation classification, and protein-level ranking. This generalization extended beyond direct stability measurements, as OmegaTherm predictions also recovered mutation-induced stability fitness landscapes measured by DMS and enriched high-fitness variants across different Top-k selection ranges. These results support the central premise of unified stability modeling: thermodynamic and thermal stability measurements can provide complementary supervision for a shared representation of mutation responses while retaining their measurement-specific characteristics.

Ablation analyses attributed these results to the major design principles of OmegaTherm. Removing sequence-derived contact information consistently reduced performance for both ΔΔG and ΔT_m_, supporting the contribution of inferred structural context. Mutation-supervised finetuning substantially outperformed frozen pretrained representations, indicating that general evolutionary representations require adaptation to resolve subtle physical perturbations between wild-type and mutant states. Distillation improved learning from limited experimental supervision, whereas joint training improved prediction performance and reproduced the experimentally observed relationship between ΔΔG and ΔT_m_. Together, these results provide mechanistic support for the three design principles and further support unified modeling of complementary stability responses.

Despite these encouraging results, several limitations remain. First, the limited size and experimental variability of available ΔT_m_ datasets continue to constrain model performance despite the benefits of cross-measurement learning. Second, OmegaTherm primarily represents mutations through sequence-derived residue and contacts information, without explicitly modeling side-chain conformational rearrangements or fine-grained structural energetics. Incorporating side-chain geometry, physicochemical interaction potentials, or experimentally resolved structural information may further improve the modeling of mutation-induced structural perturbations. Third, OmegaTherm fundamentally transfers evolutionary information from a pretrained PLM into mutation-centered physical representations. Although the PLM encoder is fully finetuned, its initialization determines the evolutionary and structural information available for subsequent adaptation, and model performance may therefore remain dependent on the choice and quality of the underlying pretrained model.

Overall, our study suggests that mutation-induced protein stability responses should be viewed not as a collection of independent prediction tasks, but as a unified representation learning problem across heterogeneous biochemical measurements. Extending this paradigm beyond ΔΔG and ΔT_m_ to additional measurements, such as protein fitness, enzymatic activity, solubility, binding affinity, and expression level, may enable the development of more comprehensive representations of protein mutational landscapes. Such unified representations could provide a general machine intelligence framework for understanding protein sequence-function relationships and accelerating AI-guided protein engineering.

## Methods

### Datasets

All mutation data were collected from ProThermDB^45^, FireProtDB^46^ and ThermoMutDB^47^. These records were used to construct four training datasets: S8754 for ΔΔG single mutations^18^, M2339 for ΔΔG multiple mutations, S4346 for ΔT_m_ single mutations^18^ and M718 for ΔT_m_ multiple mutations. For ΔΔG evaluation, the single mutation test dataset was the independent benchmark S461^11^, and the multiple mutation test dataset was M223, which was extracted from the same source databases after redundancy filtering. For ΔT_m_ evaluation, the single and multiple mutation test datasets were S571^18^ and M286, respectively. To avoid sequence-level information leakage, all test proteins were filtered to have no more than 25% sequence identity to proteins in the corresponding training datasets. In addition, to enable a fair comparison between OmegaTherm and GeoDDG on deep mutational scanning (DMS) data, we used a sampled DMS benchmark containing 20,000 mutations from 73 proteins with available native structures^42^.

### PLM encoder

To learn evolutionary and conformational information of protein sequence, OmegaTherm uses the widely adopted protein language model ESM2 as its encoder. The pretrained ESM2 weights were used as initial parameters. The encoder consists of a token embedding layer, 33 stacked transformer layers and a linear regression head for contact map prediction (Supplementary Fig. 9). Given a wild-type or mutant protein sequence p = {r_1_, r_2_, . . ., r_L_} with L residues, where r_i_ ∈ {A, C, . . ., Y} denotes the residue type at position *i*, the sequence was first converted into tokens:

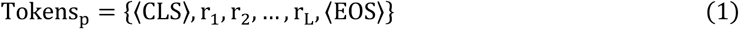

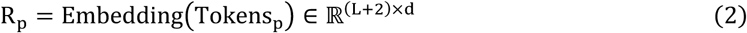

where Tokens_p_ denotes the tokenized protein sequence, ⟨CLS⟩ and ⟨EOS⟩ are two additional tokens that indicate the beginning and end of the sequence, and d is the representation dimension. The token representations R_p_ were then passed through the transformer layers:

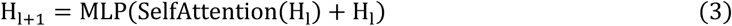

where H_l_ ∈ ℝ^(L+2)×d^ is the input of (l + 1) -th transformer layer (l = 0, . . ., n − 1), and H_0_ = R_p_ initially. SelfAttention(·) denotes the standard attention mechanism that generates query, key and value matrices, calculates attention scores between tokens and updates their representations. MLP(·) is composed of several linear layers and corresponding nonlinear activate functions. Through transformer layers, attention maps from different layers and heads were concatenated to predict residue-residue contact probabilities:

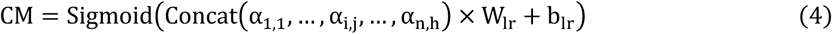

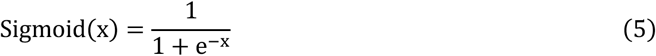

where α_i,j_ is the attention map from the j -th attention heads in i -th transformer layer, (W_lr_ ∈ ℝ^(n×h,1)^, b_lr_ ∈ ℝ^1^) is a linear layer, and (n, h) represent the number of transformer layers and attention heads per layer, respectively.

### Mutation-pair construction module

The PLM encoder generated residue representations H_n_and corresponding contact map CM for each target protein sequence p. Based on these outputs, OmegaTherm introduces a mutation-pair construction module to generate neighbor representations for the mutation sites {m_1_, m_2_, … } (1 ≤ m_i_ ≤ L).

For each mutation site, OmegaTherm first selected sequential and spatial neighbors. Sequential neighbors were defined as the two residues immediately upstream and downstream of the mutation site. Spatial neighbors were selected as the top-k residues with the highest predicted contact probabilities:

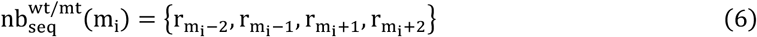

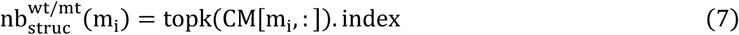

where k is set as 20. The neighbor sets from the wild-type and mutant sequences were then combined as the final selected residues:

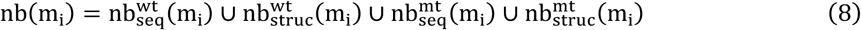

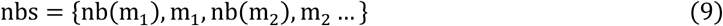

The retained residues were used to construct paired features from the wild-type and mutant PLM embeddings. For each mutation site m_i_, OmegaTherm concatenated the wild-type embedding, mutant embedding, their difference and their element-wise product, and then aggregated them into a paired feature matrix, denoted as *Pair* ∈ ℝ^|*nbs*|×*d*^:

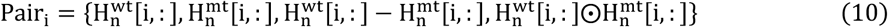

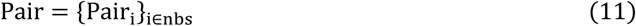

### Measurement-specific mutation-centered encoder

The measurement-specific mutation-centered encoder aggregates paired representations of retained residues into a mutation-level representation. To describe the relationship between each retained residue and the mutation sites, we constructed two types of discrete information: residue-neighbor type and relative distance to mutation sites.

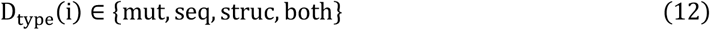

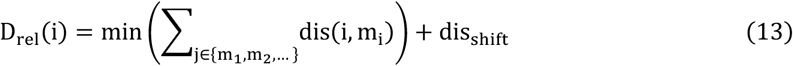

where i is a retained residue and dis_shift_is set as 32 to ensure that all position values are non-negative. The four residue-neighbor types, mut, seq, struc and both, indicate a mutation site, a sequential neighbor, a spatial neighbor and an overlapping sequential-spatial neighbor, respectively. D_rel_(i) is the binned shortest sequence-level distance from residue i to the known mutation sites {m_1_, m_2_, … }. These discrete features were embedded and added to the paired residue representations:

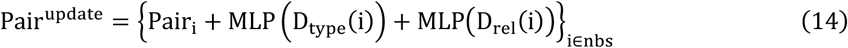

Subsequently, an additional ⟨CLS⟩ token was prepended to the paired residue representations Pair^update^. The resulting sequence was processed by two self-attention and feed-forward layers. Finally, the updated ⟨CLS⟩ representation was concatenated with environmental factors and used to predict ΔΔG and ΔT_m_ . The ΔΔG environmental factors included pH and experimental temperature, whereas the ΔT_m_ contained pH only.

### Distillation strategy and model training

As shown in Fig. 1e, self-distillation consisted of three stages. In the first stage, two models were trained for each task: a lightweight PLM model (Supplementary Figs. 10 and 11) and OmegaTherm. In this stage, the ESM block was fixed, and the remaining model components were trained. In the second stage, the full model was unfrozen and finetuned. The trained models were then applied to the training data from the other task, and the average of their predictions was used as the pseudo labels. These data with pseudo labels were used to further finetune the previous OmegaTherm model in the third stage. In parallel, the combined data and the stage-1 pretrained OmegaTherm models were used to train the unified student model (Fig. 1f).

To filter out unreliable pseudo labels, we defined a confidence range based on experimentally validated paired data in the training set (Supplementary Fig. 12). For each mutation sample, pseudo label reliability was assessed using the relationship between its true label in one task and its pseudo label in the other task. The filter criteria were as follows:

1. The pseudo label was required to have the same sign as the true label from the other task (e.g., sign(ΔΔG_pseudo_H = sign xΔT_mexp_)≤θ).
2. The pseudo-label value had to fall within an acceptable range relative to the scaled experimental label (e.g., abs (ΔΔG_pseudo_ − s × ΔT_mexp_) ≤ θ). where *s* is a scaling parameter estimated from known paired training data (Supplementary Fig. S11), θ = 0.5 + 0.25 × max (abs(ΔΔG_pseudo_), abs (*s* × ΔT_mexp_)) is a threshold determined jointly by experimental values and pseudo label values.

To reduce the influence of outlier samples, OmegaTherm was trained using a smooth L1 loss. A simple ranking loss was also used in the first stage to improve mutation-ranking performance:

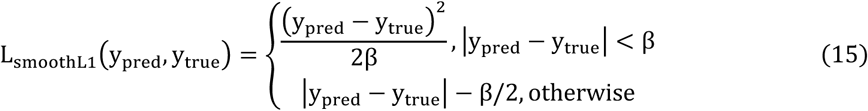

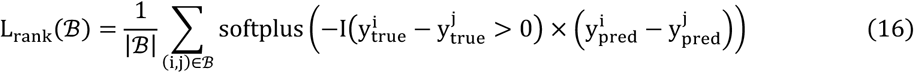

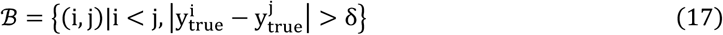

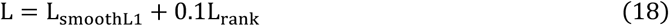

where β is set as 1 to reduce the impact of outliers and ℬ consists of sample pairs in the batch whose labels differ by a certain margin δ. L_rank_(ℬ) is used to improve the model’s ability to rank these sample pairs in same batches. Single and multiple mutation samples were jointly trained within each assay.

For a single mutation, the mutation-centered encoder received features associated with one mutation site. For multiple mutations, it received a variable-length collection of site-associated features and aggregated information across the mutated sites.

All experiments of OmegaTherm are carried out using one NVIDIA Tesla V100s GPU card with 32 GB of memory. The batch size for stage 1 training is 128 while it is 1 for all other stages. Training took approximately 16 hours.

### Mutation-sensitive site detection

To characterize mutation sensitivity, OmegaTherm was used to predict all possible single residue substitutions in a target protein. These predictions were used to construct a single mutation landscape and identify positions at which substitutions were more likely to destabilize the protein. For each position pos_i_, we calculated a sensitivity score as the mean destabilizing effect across all non-wild-type amino acid substitutions:

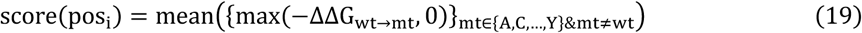

A higher score(pos_i_) indicates that substitutions at position i are more likely to destabilize the protein, and therefore that the position is more sensitive. After sensitive sites were detected, residues within a two-residue window on either side of each sensitive site were defined as part of the corresponding sensitive region.

### Evaluation metrics

Model performance was evaluated using regression and multiclass classification metrics. For regression, we used Spearman correlation coefficient (SCC), Pearson correlation coefficient (PCC), coefficient of determination (R^2^), root mean square error (RMSE), mean absolute error (MAE) and median absolute error (MedAE). For classification, we used accuracy (ACC), Matthews correlation coefficient (MCC) and Cohen’s kappa score. Specifically, given predicted results *Y_pred_* and true labels *Y_true_*, SCC is defined as:

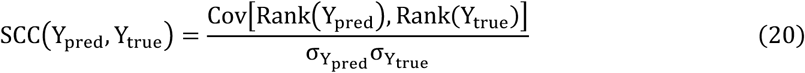

where Rank(·) is the rank positions of elements in the sorted list. Cov denotes the covariance between two lists, and *σ* is the standard deviation. PCC was calculated directly from the predicted and true values:

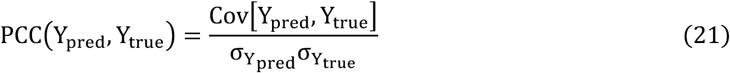

The coefficient of determination was calculated as:

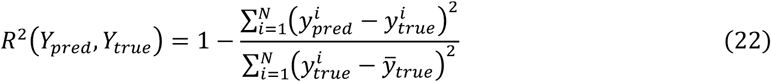

where larger SCC, PCC and R^2^ values indicate better predictive performance. RMSE and MAE were used to measure prediction errors:

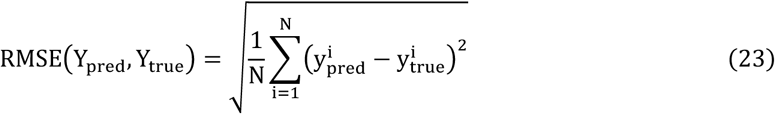

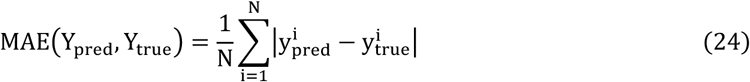

Both smaller RMSE and MAE indicate smaller prediction errors. Additional three metrics, ACC, MCC, and Cohen’s kappa score (*κ*) evaluate the multi-class classification abilities of models:

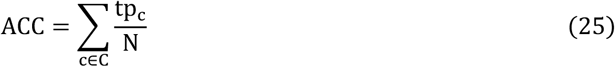

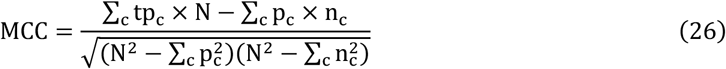

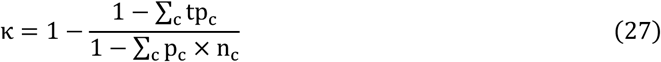

where *tp_c_* denotes the number of true positives for category *c*, and *N* represents the number of samples. *p_c_* and *n_c_* represent the number of samples predicted as class *c* and the number of samples belonging to class *c*, respectively. Both higher ACC, MCC, and *κ* values show better performance in multi-class classification.

## Data and Software Availability

The data and source code can be downloaded from https://github.com/CSUBioGroup/OmegaTherm .

## Acknowledgments

This work is supported in part by the National Natural Science Foundation of China under Grant (No.625B2184 to W.W. and No.62225209 to M.L.), Singapore Ministry of Education (T1 251RES2309 to Y.Z.), National University of Singapore Startup Grants (#A-8001129-00-00 and #A-0009651-30-00 to Y.Z.), the National Key Research and Development Program of China (2023YFF1205500 to R.Z.), and the High Performance Computing Center of Central South University. The authors thank Prof. Nam-Hai Chua and his team for stimulating discussions which helped initiate this project.

## Author Contributions

W.W., X.H., M.L., and Y.Zhang conceived and designed the study; W.W. and Y.Zhou designed the model and methodology; W.W. implemented the model and performed the computational experiments; W.W. and X.H. analyzed data; W.W., Y.Zhou, and W.Y. collected and constructed benchmark datasets; Q.Y. and F.X. contributed to data visualization and result interpretation. X.H., Y.W., F.X., and R.Z. contributed to the methodological discussions and experimental design; W.W., Y.Zhou, W.Y., Y.W., Q.Y., F.X., X.H., M.L., and Y.Zhang wrote and revised the paper.

