## Supplementary Files for "Unified modeling of mutation-induced protein stability responses with OmegaTherm"

### with OmegaTherm

Wenkang Wang<sup>a,b†</sup>, Yihang Zhou<sup>b,ct</sup>, Wentao Yu<sup>a</sup>, Yifan Wu<sup>a,c</sup>, Qiurong Yang<sup>a</sup>, Fanding Xu<sup>b,d</sup>, Ruiqing Zheng<sup>a</sup>,
Xiaoqiang Huang<sup>e\*</sup>, Min Li<sup>a\*</sup>, Yang Zhang<sup>b,c,f\*</sup>

<sup>a</sup> School of Computer Science and Engineering, Central South University, Changsha, 410083, China

<sup>b</sup> Department of Computer Science, School of Computing, National University of Singapore, Singapore, 117417, Singapore

<sup>c</sup> Cancer Science Institute of Singapore, National University of Singapore, Singapore, 117599, Singapore

<sup>d</sup> School of Life Science and Technology, Xi'an Jiao Tong University, Shanxi, 710049, China

<sup>e</sup> Center for Advanced Models for Translational Sciences and Therapeutics, Department of Internal Medicine, University of Michigan, Ann Arbor, MI 48109, USA

<sup>f</sup> Department of Biochemistry, Yong Loo Lin School of Medicine, National University of Singapore, Singapore, 117596, Singapore

<sup>†</sup> W.W. and Y.Zhou contributed equally to this work

This PDF file includes:

Supplementary Table 1 to 6.

Supplementary Figure 1 to 12.

**Supplementary Table 1.** Overall comparison of $\Delta\Delta G$ prediction on S461 in terms of SCC, PCC, RMSE, and MAE.

| Method | Total |  | Direct |  | Inverse |  | Antisymmetry |  |
| --- | --- | --- | --- | --- | --- | --- | --- | --- |
| | PCC/SCC/R2 | RMSE/MAE | PCC/SCC/R2 | RMSE/MAE | PCC/SCC/R2 | RMSE/MAE | $r$ | $\delta$ |
| *Sequence-based* |  |  |  |  |  |  |  |  |
| **OmegaTherm** | **0.83/0.85/0.68** | **0.95/0.67** | 0.68/**0.69/0.42** | **0.95/0.67** | 0.68/**0.69/0.42** | **0.95/0.67** | **-1.00** | **0.00** |
| **DDGEmb** | **0.83/0.83/0.67** | 0.96/0.74 | 0.67/0.66/0.40 | 0.97/0.74 | 0.67/0.67/0.41 | 0.96/0.74 | -0.98 | 0.11 |
| **I-Mutant3.0-Seq<sup>a</sup>** | 0.47/0.45/0.00 | 1.69/1.32 | 0.44/0.45/0.14 | 1.16/0.89 | 0.24/0.25/-1.78 | 2.09/1.75 | -0.43 | -1.57 |
| **SAAFEC-Seq<sup>a</sup>** | 0.35/0.25/-0.13 | 1.80/1.37 | 0.49/0.48/0.20 | 1.12/0.83 | 0.02/0.01/-2.33 | 2.29/1.92 | -0.05 | -1.70 |
| **DDGun<sup>a</sup>** | 0.76/0.77/0.44 | 1.29/0.96 | 0.59/0.58/-0.00 | 1.26/0.95 | 0.55/0.55/-0.09 | 1.32/0.97 | -0.94 | -0.14 |
| **INPS-Seq<sup>a</sup>** | 0.77/0.79/0.57 | 1.11/0.82 | 0.56/0.55/0.23 | 1.10/0.82 | 0.55/0.54/0.21 | 1.12/0.83 | **-1.00** | -0.01 |
| **ACDC-NN-Seq<sup>a</sup>** | 0.76/0.76/0.58 | 1.10/0.80 | 0.57/0.54/0.23 | 1.10/0.80 | 0.57/0.54/0.23 | 1.10/0.80 | **-1.00** | **0.00** |
| **MUPro<sup>a</sup>** | 0.47/0.43/-0.11 | 1.79/1.42 | 0.40/0.37/0.13 | 1.17/0.91 | 0.29/0.27/-2.20 | 2.24/1.93 | -0.31 | -1.93 |
| *Structure-based* |  |  |  |  |  |  |  |  |
| **GeoDDG-AF2<sup>b</sup>** | 0.82/0.83/0.66 | 0.98/0.73 | **0.69/0.68/0.38** | 0.98/0.73 | **0.69/0.68/0.38** | 0.98/0.73 | **-1.00** | **0.00** |
| **GeoDDG-3D<sup>b</sup>** | 0.81/0.83/0.64 | 1.01/0.72 | 0.67/0.67/0.35 | 1.01/0.72 | 0.67/0.67/0.35 | 1.01/0.72 | **-1.00** | **0.00** |
| **I-Mutant3.0<sup>a</sup>** | 0.41/0.34/-0.05 | 1.74/1.33 | 0.49/0.48/0.20 | 1.12/0.83 | 0.13/0.11/-2.05 | 2.19/1.83 | -0.04 | -1.61 |
| **FoldX<sup>a</sup>** | 0.44/0.53/-0.40 | 2.00/1.26 | 0.22/0.35/-2.18 | 2.23/1.38 | 0.41/0.45/-0.94 | 1.74/1.14 | -0.22 | -0.45 |
| **SDM<sup>a</sup>** | 0.41/0.33/0.01 | 1.69/1.26 | 0.56/0.51/-0.13 | 1.33/1.01 | 0.11/0.11/-1.50 | 1.98/1.51 | -0.35 | -0.84 |
| **DDGun3D<sup>a</sup>** | 0.76/0.72/0.55 | 1.14/0.83 | 0.64/0.58/0.22 | 1.10/0.80 | 0.59/0.55/0.12 | 1.17/0.85 | -0.96 | -0.13 |
| **MAESTRO<sup>a</sup>** | 0.54/0.48/0.17 | 1.54/1.16 | 0.63/0.60/0.31 | 1.04/0.78 | 0.20/0.17/-1.35 | 1.92/1.53 | -0.22 | -1.18 |
| **mCSM<sup>a</sup>** | 0.46/0.40/-0.02 | 1.71/1.31 | 0.53/0.51/0.27 | 1.07/0.80 | 0.18/0.12/-2.00 | 2.17/1.81 | -0.09 | -1.64 |
| **ACDC-NN<sup>a</sup>** | 0.78/0.78/0.60 | 1.07/0.77 | 0.61/0.57/0.29 | 1.06/0.77 | 0.60/0.56/0.26 | 1.07/0.78 | -0.98 | -0.04 |
| **Dynamut<sup>a</sup>** | 0.63/0.59/0.40 | 1.32/0.98 | 0.50/0.44/-0.02 | 1.27/0.96 | 0.44/0.43/-0.19 | 1.36/1.01 | -0.61 | -0.11 |
| **INPS3D<sup>a</sup>** | 0.69/0.71/0.44 | 1.27/0.93 | 0.62/0.60/0.35 | 1.01/0.75 | 0.35/0.39/-0.39 | 1.48/1.12 | -0.49 | -0.67 |
| **PopMusic<sup>a</sup>** | 0.60/0.56/0.20 | 1.51/1.14 | 0.62/0.61/0.34 | 1.01/0.76 | 0.29/0.25/-1.27 | 1.89/1.52 | -0.30 | -1.34 |
| **ThermoNet<sup>a</sup>** | 0.67/0.60/0.44 | 1.27/0.97 | 0.56/0.49/0.04 | 1.23/0.93 | 0.52/0.43/-0.10 | 1.31/1.01 | -0.86 | -0.11 |
| **DUET<sup>a</sup>** | 0.52/0.46/0.12 | 1.58/1.18 | 0.59/0.57/0.28 | 1.06/0.77 | 0.18/0.15/-1.49 | 1.97/1.59 | -0.15 | -1.29 |
| **PremPS<sup>a</sup>** | 0.81/0.81/0.64 | 1.01/0.76 | 0.63/0.60/0.32 | 1.03/0.79 | 0.63/0.58/0.38 | 0.98/0.73 | -0.86 | 0.23 |

<sup>a</sup> indicates that the raw results are taken from a prior benchmark. <sup>b</sup> refers that the raw results are taken from its original paper.

**Supplementary Table 2.** Statistical analysis on S461 and M223

| Data | Method | Median Absolute Error | P-value |
| --- | --- | --- | --- |
| S461 | OmegaTherm | 0.42 |  |
|  | DDGEmb | 0.58 | 1.59e-05 |
|  | GeoDDG-3D | 0.51 | 1.90e-05 |
|  | PremPS | 0.60 | 1.91e-07 |
|  | INPS3D | 0.69 | 4.36e-26 |
| M223 | OmegaTherm | 0.69 |  |
|  | DDGEmb | 1.17 | 1.57e-13 |
|  | GeoDDG-AF2 | 1.75 | 5.28e-39 |

**Supplementary Table 3.** Overall comparison of $\Delta\Delta G$ prediction on M223 in terms of SCC, PCC, RMSE and MAE.

| Method | Total |  | Direct |  | Inverse |  | Antisymmetry |  |
| --- | --- | --- | --- | --- | --- | --- | --- | --- |
| | PCC/SCC/R2 | RMSE/MAE | PCC/SCC/R2 | RMSE/MAE | PCC/SCC/R2 | RMSE/MAE | r | $\delta$ |
| **OmegaTherm** | **0.77/0.84/0.56** | **2.07/1.16** | **0.77/0.82/0.56** | **2.07/1.16** | **0.77/0.82/0.56** | **2.07/1.16** | **-1.00** | **-0.00** |
| **DDGEmb** | 0.68/0.75/0.36 | 2.49/1.58 | 0.67/0.66/0.35 | 2.51/1.61 | 0.68/0.70/0.36 | 2.48/1.55 | -0.97 | -0.06 |
| **GeoDDG-AF2** | 0.29/0.28/-0.19 | 3.41/2.45 | 0.29/0.30/-0.18 | 3.38/2.43 | 0.28/0.28/-0.22 | 3.43/2.46 | -0.99 | 0.04 |

**Supplementary Table 4.** Overall comparison of $\Delta T_m$ prediction on S571 in terms of SCC, PCC, RMSE, and MAE.

| Method | Total |  | Direct |  | Inverse |  | Antisymmetry |  |
| --- | --- | --- | --- | --- | --- | --- | --- | --- |
| | PCC/SCC/R2 | RMSE/MAE | PCC/SCC/R2 | RMSE/MAE | PCC/SCC/R2 | RMSE/MAE | r | $\delta$ |
| *Sequence-based* |  |  |  |  |  |  |  |  |
| **OmegaTherm** | **0.64/0.60/0.39** | **7.41/4.89** | **0.59/0.58/0.30** | **7.41/4.89** | **0.59/0.58/0.30** | **7.41/4.89** | **-1.00** | **0.00** |
| **Seq2Topt** | 0.30/0.20/0.09 | 9.05/6.18 | 0.17/0.04/-0.04 | 9.05/6.18 | 0.17/0.04/-0.04 | 9.05/6.18 | **-1.00** | **-0.00** |
| *Structure-based* |  |  |  |  |  |  |  |  |
| **GeoDTm-AF2<sup>a</sup>** | 0.52/0.54/0.27 | 8.11/5.55 | 0.46/0.51/0.16 | 8.11/5.55 | 0.46/0.51/0.16 | 8.11/5.55 | **-1.00** | **0.00** |
| **GeoDTm-3D<sup>a</sup>** | 0.53/0.55/0.28 | 8.03/5.31 | 0.47/0.51/0.18 | 8.03/5.31 | 0.47/0.51/0.18 | 8.03/5.31 | **-1.00** | **-0.00** |
| **AUTO-MUTE<sup>a</sup>** | 0.29/0.25/0.08 | 8.50/5.79 | 0.29/0.25/0.08 | 8.50/5.79 | - | - | - | - |
| **HoTMuSiC<sup>a</sup>** | 0.33/0.30/0.10 | 8.41/5.70 | 0.33/0.30/0.10 | 8.41/5.70 | - | - | - | - |

<sup>a</sup> refers that the raw results are taken from a prior benchmark.

**Supplementary Table 5.** Statistical analysis on S571 and M286

| Data | Method | Median Absolute Error | P-value |
| --- | --- | --- | --- |
| S571 | OmegaTherm | 2.99 |  |
|  | GeoDTm-3D | 3.34 | 2.35e-4 |
|  | GeoDTm-AF2 | 3.70 | 7.59e-11 |
|  | HoTMuSiC | 3.86 | 9.64e-07 |
|  | AUTO-MUTE | 3.96 | 7.46e-08 |
|  | Seq2Topt | 4.03 | 6.33e-25 |
| M286 | OmegaTherm | 7.24 |  |
|  | Seq2Topt | 8.32 | 2.73e-07 |
|  | GeoDTm-AF2 | 8.54 | 8.20e-19 |

**Supplementary Table S6.** Overall comparison of $\Delta T_m$ prediction on M286 in terms of SCC, PCC, RMSE, and MAE.

| Method | Total |  | Direct |  | Inverse |  | Antisymmetry |  |
| --- | --- | --- | --- | --- | --- | --- | --- | --- |
| | PCC/SCC/R2 | RMSE/MAE | PCC/SCC/R2 | RMSE/MAE | PCC/SCC/R2 | RMSE/MAE | r | $\delta$ |
| OmegaTherm | **0.61**/0.30/**0.34** | **10.53**/**8.44** | **0.59**/0.18/**0.30** | **10.53**/**8.44** | **0.59**/0.18/**0.30** | **10.53**/**8.44** | **-1.00** | **0.00** |
| Seq2Topt | 0.48/**0.43**/0.11 | 12.20/9.87 | 0.57/**0.49**/0.06 | 12.20/9.87 | 0.57/**0.49**/0.06 | 12.20/9.87 | **-1.00** | **-0.00** |
| GeoDTm-AF2 | 0.36/0.12/0.02 | 12.84/10.24 | 0.38/0.08/-0.06 | 12.93/10.29 | 0.39/0.10/-0.03 | 12.76/10.19 | **-1.00** | -0.05 |

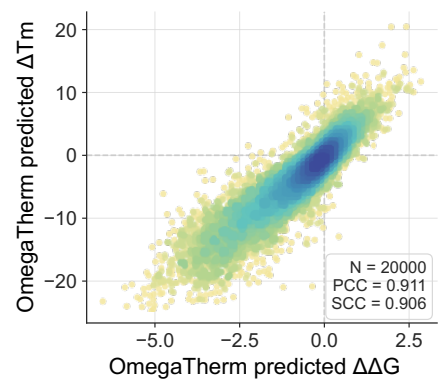

**Supplementary Figure 1.** Correlation between OmegaTherm-predicted $\Delta\Delta G$ and $\Delta T_m$ on DMS fitness dataset.

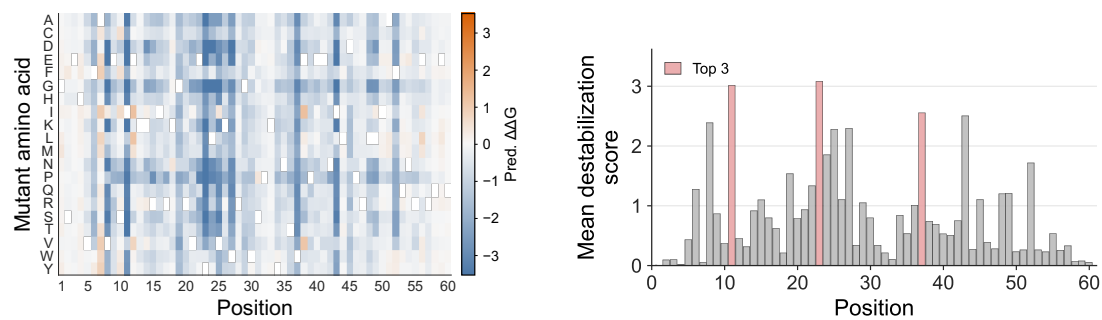

**Supplementary Figure 2.** Left panel is the complete mutation landscape of 2K28 generated by OmegaTherm. Right panel is the predicted sensitive sites of 2K28 by OmegaTherm, where a higher score indicates that mutations on this position are more likely to protein destabilization.

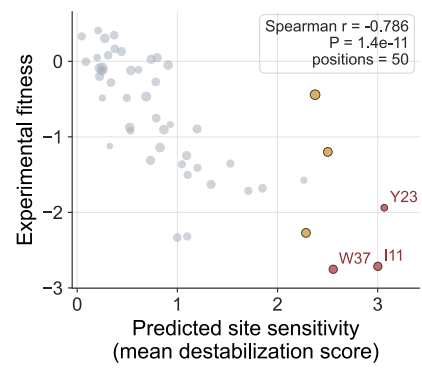

**Supplementary Figure 3.** Correlation between the predicted sensitivity ranking and the corresponding fitness variation.

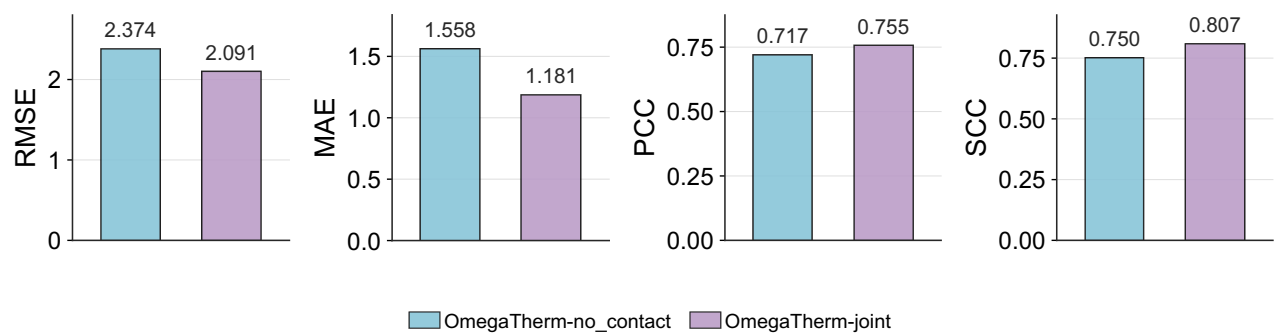

**Supplementary Figure 4.** The performance comparison of OmegaTherm and OmegaTherm w/o contact on $\Delta\Delta G$ multiple mutation dataset in terms of RMSE, MAE, PCC and SCC.

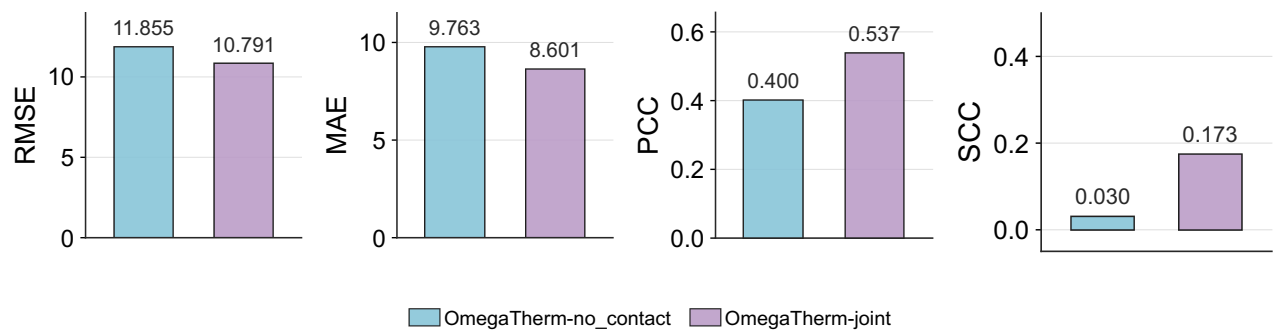

**Supplementary Figure 5.** The performance comparison of OmegaTherm and OmegaTherm w/o contact on $\Delta T_m$ multiple mutation dataset in terms of RMSE, MAE, PCC and SCC.

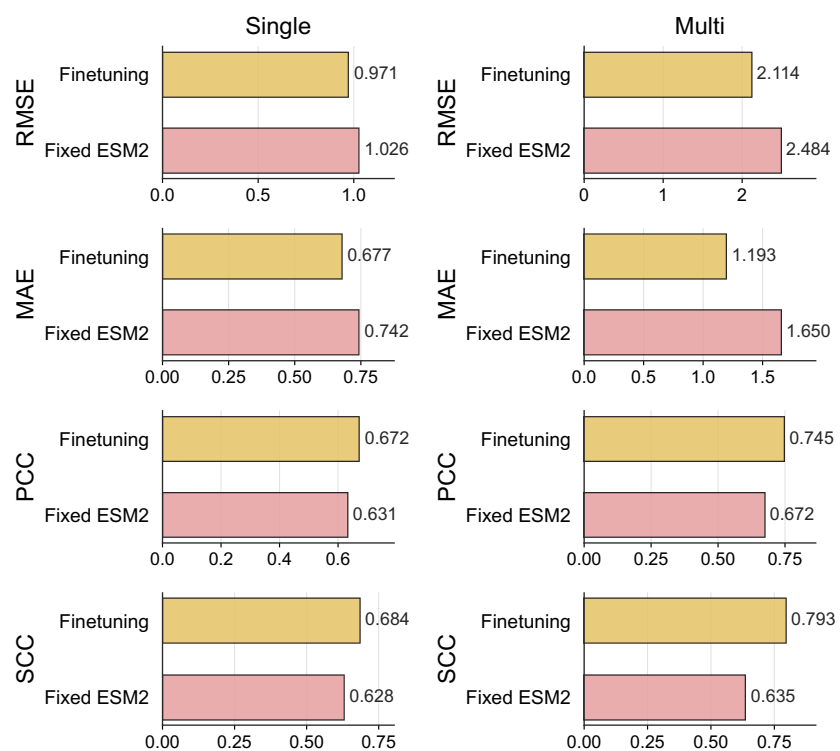

**Supplementary Figure 6.** The performance comparison between fixed ESM2 and finetuning full model on $\Delta\Delta G$ dataset.

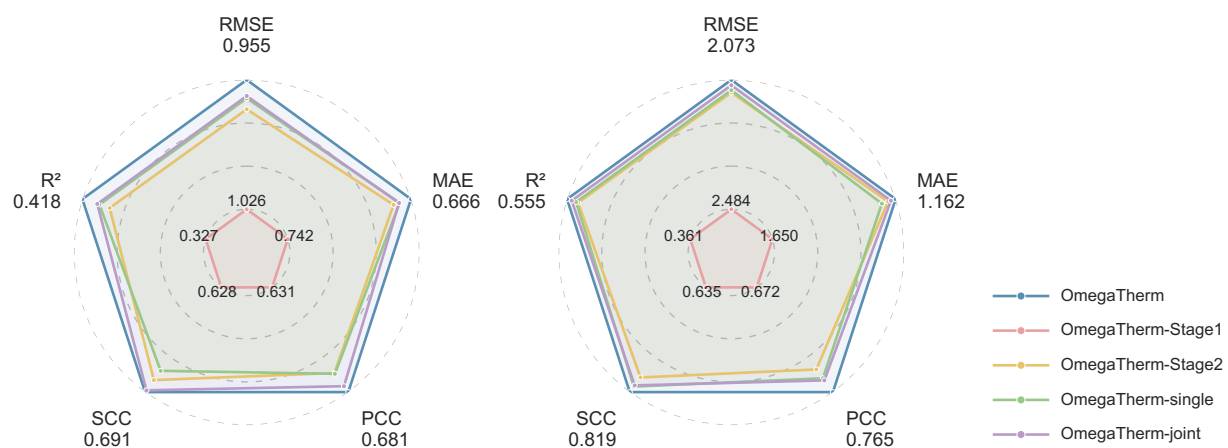

**Supplementary Figure 7.** The ablation experiment of OmegaTherm on $\Delta\Delta G$ dataset, where the left radar plot corresponds single mutations and the right radar plot denotes multiple mutations.

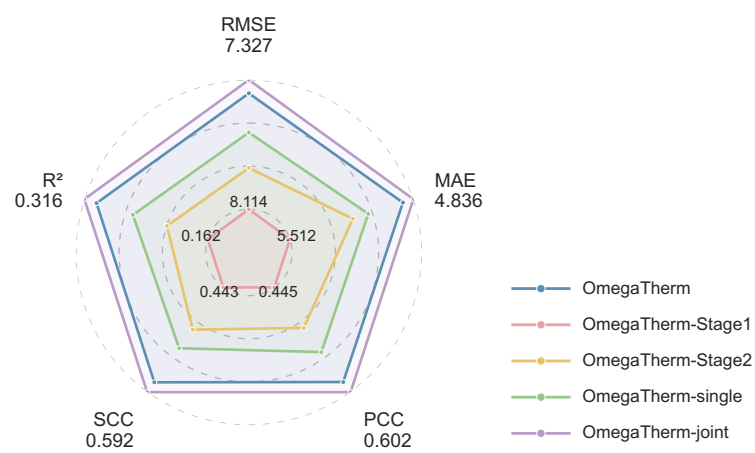

**Supplementary Figure 8.** The ablation experiment of OmegaTherm on $\Delta T_m$ single mutation dataset.

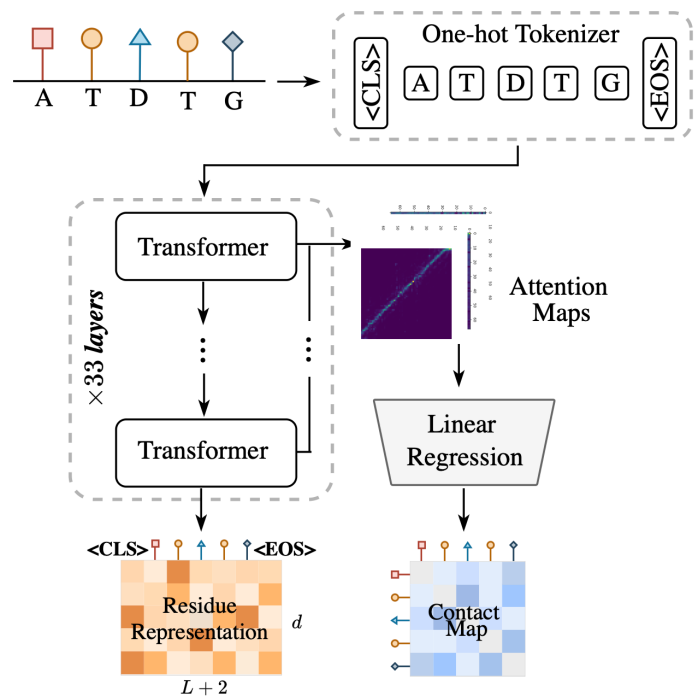

**Supplementary Figure 9.** The architecture of PLM encoder.

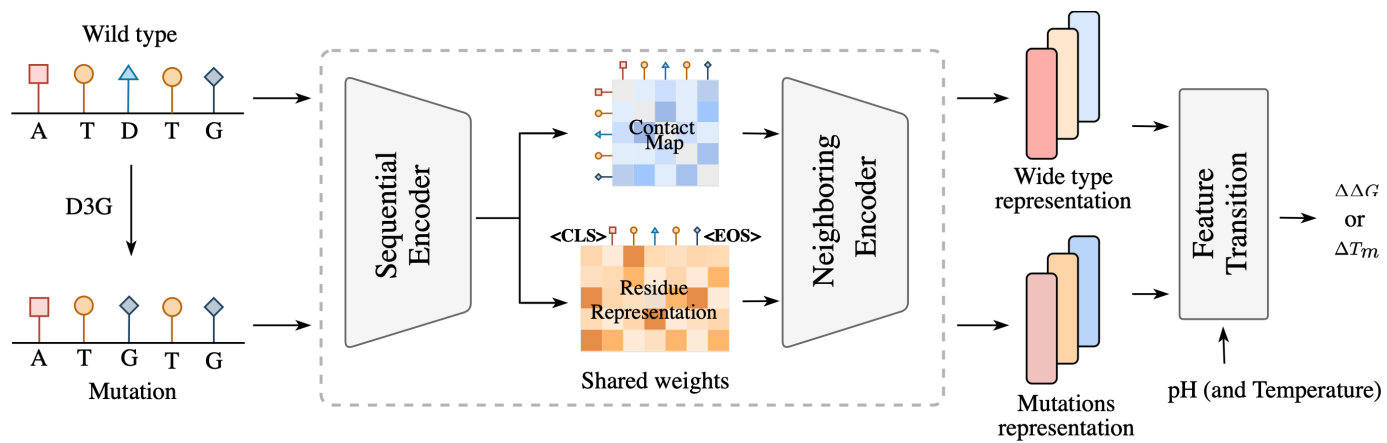

**Supplementary Figure 10.** The architecture of lightweight PLM model, consisting of a ESM encoder, a cross-attention module and a feature transition module for prediction.

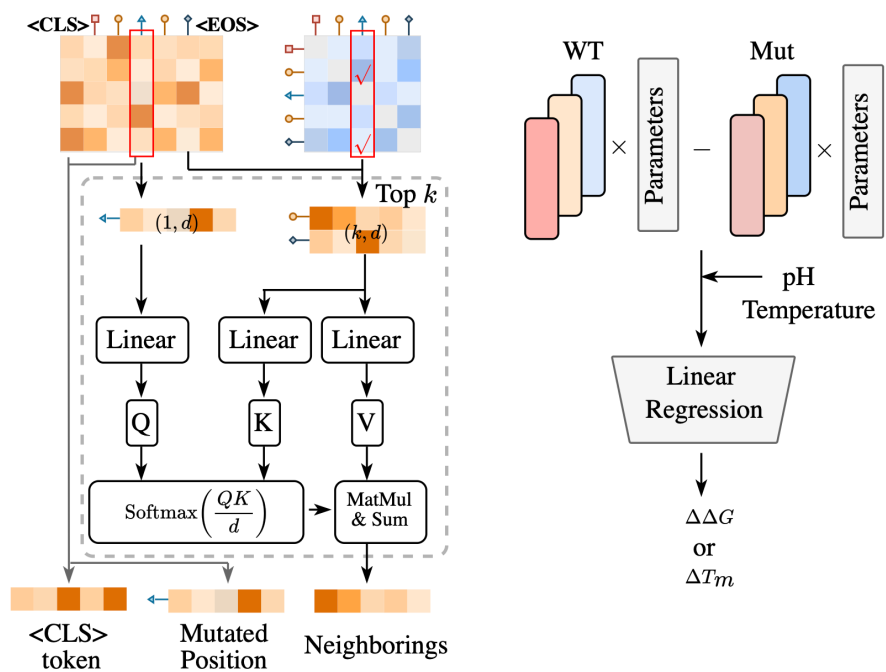

**Supplementary Figure 11.** The architecture of cross-attention module (left part) and feature transition module (right part) used in lightweight PLM model.

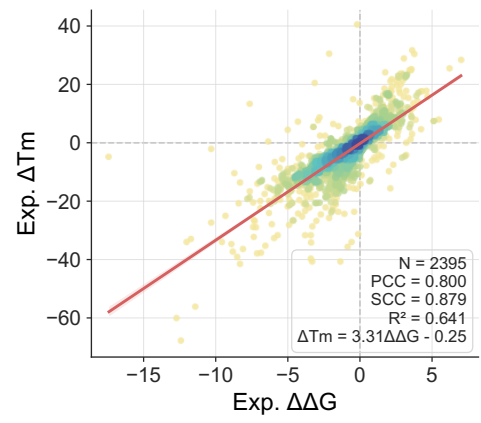

**Supplementary Figure 12.** The scaling relationship between $\Delta\Delta G$ and $\Delta T_m$.
